# Chemical suppression of a bacterial immune system revives repressed phages

**DOI:** 10.64898/2026.04.28.721336

**Authors:** Chengqian Zhang, Dziugas Sabonis, Yanyao Cai, Zhiyu Zang, Giedre Tamulaitiene, Joseph P. Gerdt

## Abstract

Many antiviral immune systems have recently been discovered in bacteria. The mechanisms of several are obscure, as are their individual significance for antiphage defense. To shed light on the mechanism and significance of the two-component type I Thoeris antiphage immune system, we leveraged high-throughput phenotypic screening to identify three small molecule inhibitors. The inhibitors target the ThsA NADase component, inhibiting its 3ʹ-cADPR-activated filamentation. The temporal control afforded by the small-molecule inhibitors allowed us to answer an outstanding question in antiviral immunity—is persistent immunity required to repress phage titers, or do immune systems become unnecessary after eradicating infectious phages? We found that Thoeris immunity must be maintained, as chemical inhibition enabled repressed phages to revive and overtake the bacterial population. Furthermore, due to the cooperative nature of antiviral immunity, we found that Thoeris must be inhibited in only 10% of the bacteria to cause phage-induced lysis of the entire population.

## Introduction

Bacteria are constantly exposed to bacteriophage (phage) infections, which pose a major threat to their survival. ^1,2^ To counteract these viral predators, bacteria have evolved a diverse collection of defense mechanisms. ^3–5^ In addition to nuclease defense systems (e.g., Restriction-Modification^6^ and CRISPR-Cas^7^), a series of defenses featuring abortive infection mechanisms has been widely reported. ^8–10^ The bacteria carrying abortive infection defense will execute cell death after infection to restrict the phage life cycle and prevent the release of propagated phages. ^11^ Although bioinformatic discovery has presented a massive bacterial immuno-weaponry, the world’s understanding of their molecular mechanisms and functional roles remains incomplete. ^5^

The development and application of chemical probes can help fill this gap in understanding. Chemical inhibitors are indispensable tools to dissect biochemical mechanisms and investigate the timing and functional roles of biological processes. ^12–14^ Moreover, the rapid global dissemination of multidrug-resistant (MDR) bacteria has revitalized interest in phage therapy as a promising alternative to conventional antibiotics. In this context, chemical inhibitors are not only chemical probes for mechanism studies, but they may also broaden the therapeutic window of phage therapies. For example, chemical inhibition of bacterial antiviral immune systems should allow phages to kill ‘phage-resistant’ pathogens. ^15^ Therefore, inhibitors of antiphage immune systems provide opportunities to develop next-generation precision antimicrobials.

In this manuscript, we aimed to discover and apply small molecule inhibitors to study the recently discovered type I Thoeris immune system. ^8^ This immune system is composed of two proteins, ThsB and ThsA (Fig. 1A). ThsB is believed to be activated by a phage protein, ^16^ after which it produces 3′-cADPR from NAD^+^.^17^ This 3′-cADPR signal molecule then binds to the SLOG domain of ThsA and triggers filamentation of ThsA. ^17–21^ It is believed that this filamentation process assembles the catalytic site of the SIR2 domain of ThsA, increasing its rate of NAD^+^ hydrolysis. ^21^ NAD^+^ depletion arrests phage replication and ultimately causes the infected cell to die in a process termed ‘abortive infection’ (Fig. 1B).

**Figure 1.**
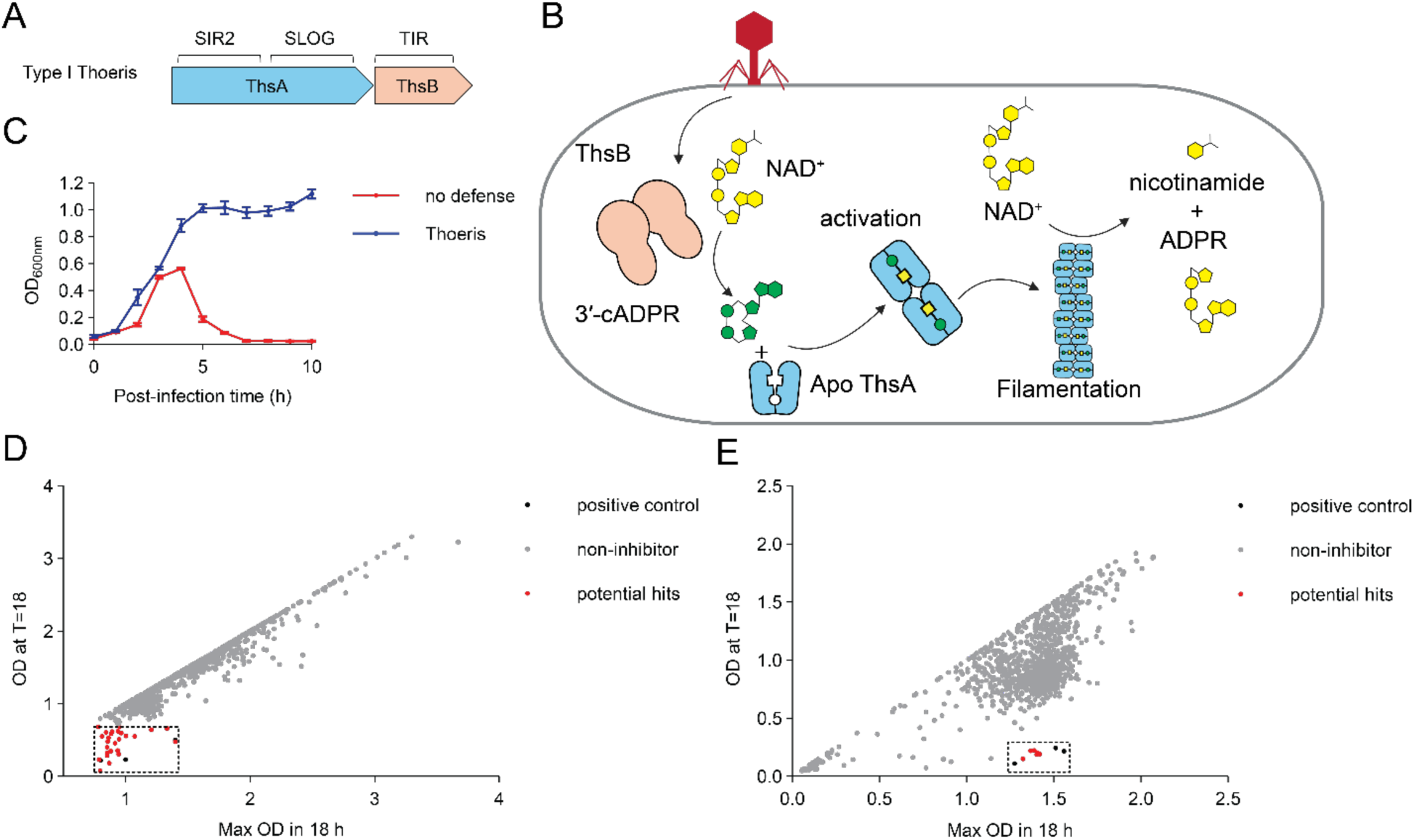
High-throughput screens identify inhibitors against type I Thoeris. A. Gene composition (domains indicated above) of a representative type I Thoeris from *B. cereus* strain MSX-D12 (SIR2, Silent Information Regulator 2 domain; SLOG, Smf/DprA-LOG domain; TIR, Toll/Interleukin-1 receptor). B. Cartoon of the protection mechanism of the type I Thoeris defense. C. Lysis curve caused by a phage cocktail against *B. subtilis* lacking immunity (red curve) compared to phage-immune *B. subtilis* possessing the Thoeris system (blue curve). Bacterial growth is monitored by optical density at 600 nm. The phage cocktail was a 1:1:1 mixture of SPO1:SP50:Goe2 at multiplicity of infection (MOI) of 0.01. Error bars represent standard error of the mean (SEM) of a biological triplicate. D–E. Screening data of the Chembridge 10,000-compound DIVERSet library (D) and the NCI 1,581-compound Diversity Set VII library (E). Both cases include three biological replicates of the ‘positive control’ (black circles), which are non-immune *B. subtilis* infected with phage. These are used to define the boundaries of putative inhibitors that inactivate Thoeris. Each grey or red circle indicates an individual tested inhibitor (grey = inactive, red = active).

We employed a high-throughput phenotypic screen of two diverse libraries of small molecules (one ∼10,000 compounds and one ∼1,500 compounds). We found inhibitors from both libraries, suggesting that even a small library is sufficient to provide useful inhibitors of antiphage immune systems. The Thoeris inhibitors resensitized phage-resistant bacteria to phages. Further in vitro experiments with purified enzymes revealed that the inhibitors prevent the 3′-cADPR-induced activation of ThsA.

Providing biochemical insight, the inhibitors validated the importance of 3′-cADPR-induced filamentation to activate the ThsA NADase. Providing functional insight, the inhibitors revealed that the Thoeris defense was not capable of eradicating phages. Instead, persistent Thoeris immunity is required to continually repress bacteriophages. Loss of immunity allows resurgence of the repressed virus. The persistence of repressed phages is conceptually related to persistent viruses in humans (e.g., the hepatitis B and C viruses), which replicate at low levels but can reactivate and spread upon immune suppression. ^22,23^ Furthermore, the existence of low levels of replicating phages suggests that inhibitor treatments alone may be sufficient to revive repressed phages in microbial communities as a ‘phage therapy’ without the need to add exogenous phages.

## Results

### High-throughput screen revealed inhibitors of a type I Thoeris system

To identify chemical inhibitors of type I Thoeris immune systems, the Thoeris operon (containing its native promoter) was cloned from *Bacillus cereus* strain MSXD12^8^ and integrated into the genome of *Bacillus subtilis* strain RM125. Plaque assays revealed that the immune system conferred protection against the SPO1, SP50, and Goe2 phages (Extended Data Fig. 1A). In order to find small molecules that can inhibit Thoeris and re-sensitize its host to phages, we set up a screen in 384-well plates containing the Thoeris-defended bacteria, a cocktail of the SPO1, SP50, and Goe2 phages (1:1:1), and synthetic compounds from screening libraries. Optical density (600 nm) of the bacterial cultures was measured every hour. Thoeris-defended bacteria grew in the presence of the phages; however, *B. subtilis* lacking Thoeris succumbed to the phage cocktail (delivered in a multiplicity of infection, MOI of 0.01) after ∼5 hours (Fig. 1C).

We screened two separate libraries of synthetic compounds: the Chembridge DIVERSet-CL Library Block 1 (10,000 compounds) (Fig. 1D) and the National Cancer Institute (NCI) Diversity Set VII (1,581 compounds) (Fig. 1E). The screening data were plotted in two dimensions, with the x-axis reporting the maximum OD_600nm_ reached during the 18-hour incubation and the y-axis reporting the final OD_600nm_ after the 18-hour incubation. Compounds that inhibit Thoeris should promote lysis curves that match the behavior of *B. subtilis* cells that lack the Thoeris defense. As shown in Fig. 1C, the final OD_600nm_ should be near zero, but the maximum OD_600nm_ should be substantially higher resulting from initial bacterial growth before the phage amplifies. Any compounds that cause near-zero readings in both axes are likely to be general antibacterials. Both screens revealed compounds with potential Thoeris inhibitory activity, shown as dots within the dashed boxes with values near the lysis curve data of the non-immune control bacteria (empty vector) with phages added (black spots).

To validate the hits, we obtained vialed compounds and repeated the lysis curves. Of the 26 potential hits tested from the Chembridge library and 6 from the NCI library, 13 and 5 (respectively) enabled reproducible lysis in a validation experiment (Fig. 2A blue, Extended Data Fig. 1B&C & Extended Data Fig. 2A&B). We then prioritized compounds that showed no cytotoxicity even at high concentrations. Among the potential inhibitors from the first round, 3 showed no toxicity in a spot-on-lawn assay (Fig. 2A green & Extended Data Fig. 2C). The 3 inhibitors were named using a library-derived prefix (CB: chembridge, NCI: NCI Diversity) followed by a serial number: CB-1, CB-6 and NCI-2 (Fig. 2B). Remarkably, CB-1 and NCI-2 showed similar structures with a urea/thiourea bridging a phenyl to an sp^3^-hybridized alkyl substructure.

**Figure 2.**
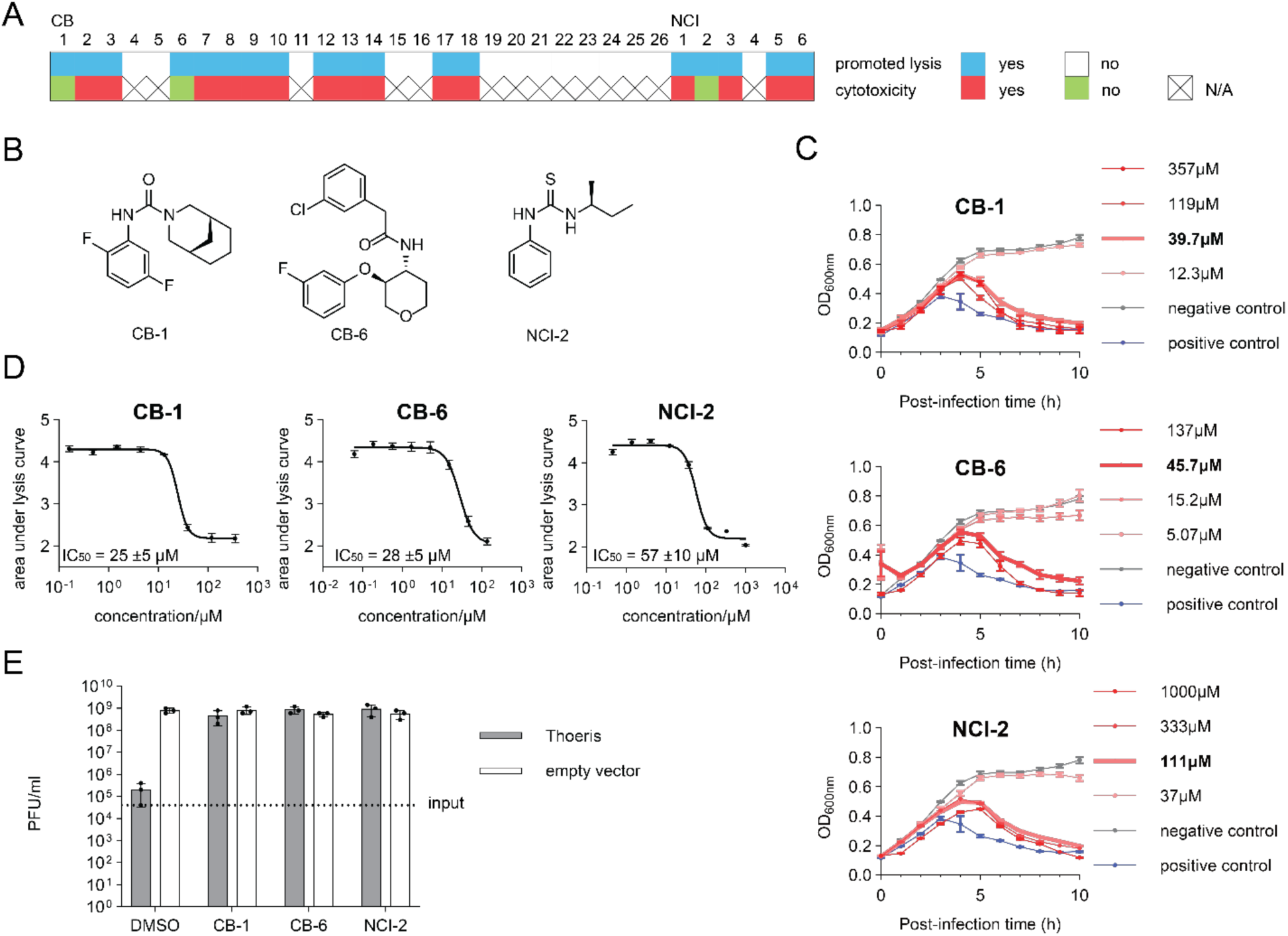
Validation of type I Thoeris inhibitors. A. Validation of potential hits from library screening. Chembridge library hits are annotated ‘CB’; NCI library hits are annotated ‘NCI’. Blue stands for type I Thoeris inhibition; white stands for no inhibition upon re-testing; red stands for cytotoxicity; green stands for no obvious cytotoxicity. Inactive compounds were not tested for toxicity (labeled “N/A”). See also Extended Data Fig. 1 and 2. B. Chemical structures of 3 reproducible non-toxic inhibitors. C. Lysis curves of type I Thoeris-expressing *B. subtilis* infected with SPO1 phage (MOI = 0.01) in the presence of different concentrations of Thoeris inhibitors. The positive control is infection of non-immune bacteria lacking Thoeris (empty vector only). The minimum inhibitory concentration (MIC) curve is bolded and labeled “(MIC)”. D. Dose-response curve of 3 inhibitors with IC_50_s displayed ± standard deviation (SD). E. Plaque-forming units (PFUs) to quantify the number of SPO1 phages that replicated on bacteria containing Thoeris or no defense (pDG) in the presence or absence of 100 µM inhibitors. In all panels, error bars represent SEM. Panels C and E have three biological replicates. Panel D has four biological replicates.

To assess their relative potencies, the inhibitors were then tested for lysis promotion at several concentrations in 96-well plates (Fig. 2C). The areas under the lysis curves were integrated and plotted as dose-response curves (Fig. 2D). All three inhibitors had similar potencies (IC_50_: 25 – 57 µM).

To further validate that these inhibitors helped the phage replicate (instead of synergizing with phage lysate to cause bacterial lysis without phage replication), we directly measured the number of infectious phage particles produced by the cultures. Indeed, all three inhibitors restored phage replication ∼10^4^× to the same level observed on host bacteria that lack the defense (Fig. 2E). Therefore, all three compounds clearly inhibit type I Thoeris, restoring phage replication and phage-induced host lysis.

### Inhibitors do not target ThsB

We next asked how each inhibitor prevents Thoeris activity. They likely either inhibit the production of the 3′-cADPR alarm signal by ThsB or the signal-induced NADase activity of ThsA. We first asked if any of the inhibitors target ThsB. To assess ThsB inhibition, we leveraged liquid chromatography mass spectrometry (LC-MS) to measure the levels of 3′-cADPR alarm signal produced by phage-infected cells. As shown in Fig. 3A, we induced expression of the ThsB protein alone in *B. subtilis*. SPO1 phage was added (MOI = 10) to activate ThsB to produce 3′-cADPR simultaneously in all cells. Cell lysates were then analyzed by LC-MS at several time points after infection, and a peak matching the m/z of 3′-cADPR (404.9987) appeared starting 40 minutes after infection (Fig. 3B). This peak’s fragmentation pattern by tandem mass spectrometry (MS/MS) matched the expected fragmentation of 3′-cADPR (Fig. 3C).

**Figure 3.**
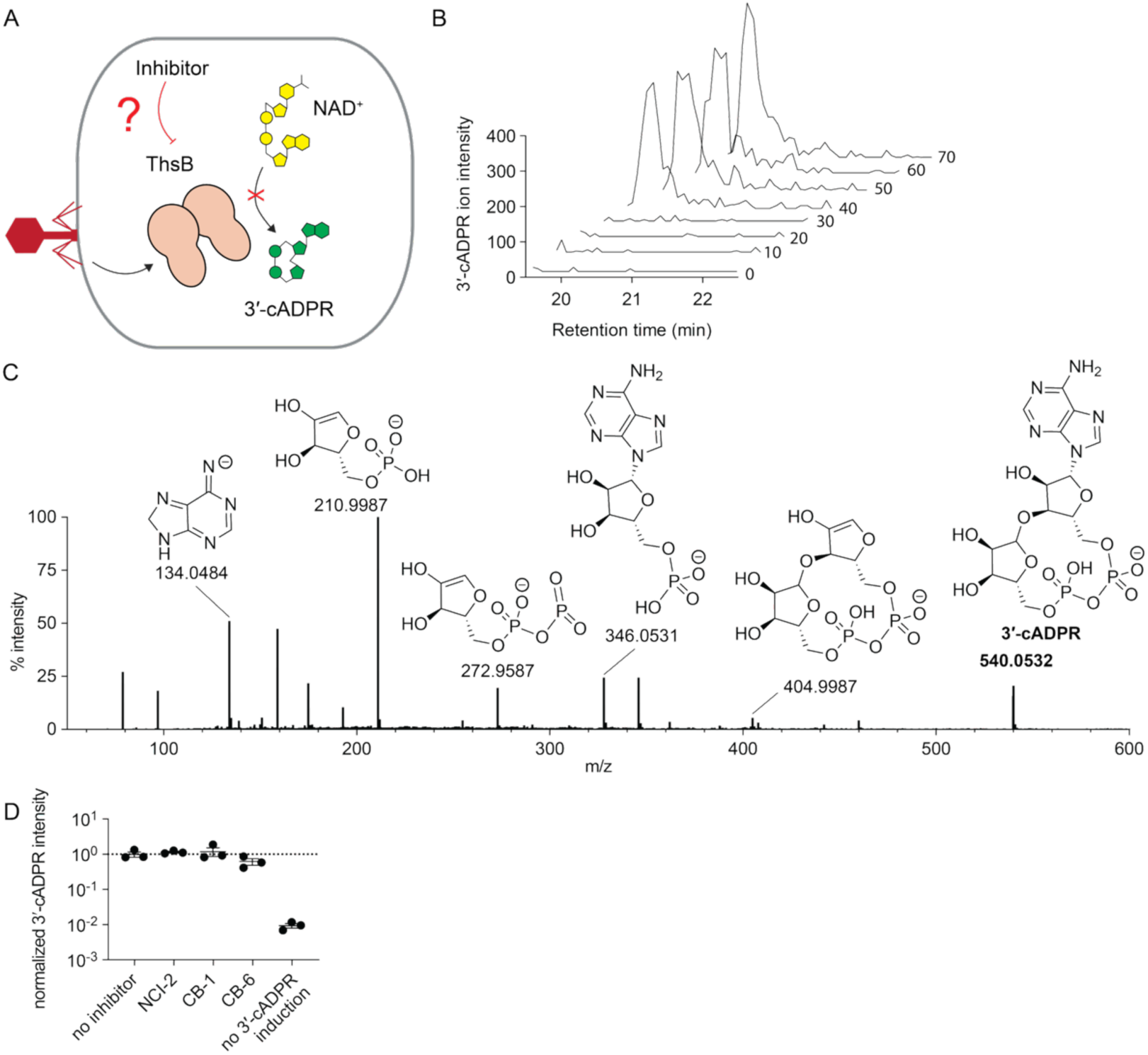
LC-MS analysis of cell lysate reveals that the inhibitors do not target ThsB. A. Schematic showing the potential effect of ThsB inhibition on 3′-cADPR signaling. B. Extracted ion chromatograms (EICs) of 3′-cADPR [M–H]^−^ (m/z 540.0424–540.0640) in the lysate of *B. subtilis* expressing ThsB 0 to 70 minutes after infection with phage SPO1 (MOI = 10). C. The MS/MS spectrum of 3′-cADPR at m/z 404.9987 shows expected fragments. Data were collected in negative ionization mode. D. Biological triplicate measurement of the normalized level of 3′-cADPR in cell lysate 50 min after infection with SPO1. 100 μM of each inhibitor was tested, and their data were normalized to DMSO-treated samples (dashed line). An uninfected “no induction” sample was used as a negative control for signal production. Data are represented as the average ± SEM from three independent biological replicates. Each replicate is displayed with a circle. One-way ANOVA with Dunnett’s multiple comparisons test (single pooled variance) showed no significant difference between any inhibitor and the ‘no inhibitor’ control (adjusted P values: NCI-2, 0.93; CB-1, 0.83; CB-6, 0.43).

Having established a method to quantify signal production by phage-activated ThsB, we tested if any of the three inhibitors blocked ThsB activity. As shown in Fig. 3D, none of the three inhibitors impacted the production of the 3′-cADPR alarm signal. Therefore, none of the inhibitors appear to target ThsB.

### Inhibitors prevent signal-induced filamentation and activation of ThsA

Since the inhibitors did not arrest alarm signal production, we hypothesized that they inhibit the other Thoeris component, ThsA. To assess ThsA inhibition, we leveraged the fluorogenic NAD^+^ analog εNAD^+^. In the presence of the 3′-cADPR signal, ThsA hydrolyzes εNAD^+^, generating a fluorescent signal in vitro. ^20,24^ We assessed if the three inhibitors could prevent this NADase activity of ThsA. Indeed, at 50 µM doses, all three inhibited εNAD^+^ hydrolysis (Fig. 4A). We assessed the potency of each inhibitor in vitro and found that they were similar (IC_50_: 18 – 54 µM, Fig. 4B), as previously observed with their potencies in sensitizing bacteria to phage infection (Fig. 2D). We noticed solubility limitations for compound CB-6, so we excluded it from further analysis.

**Figure 4.**
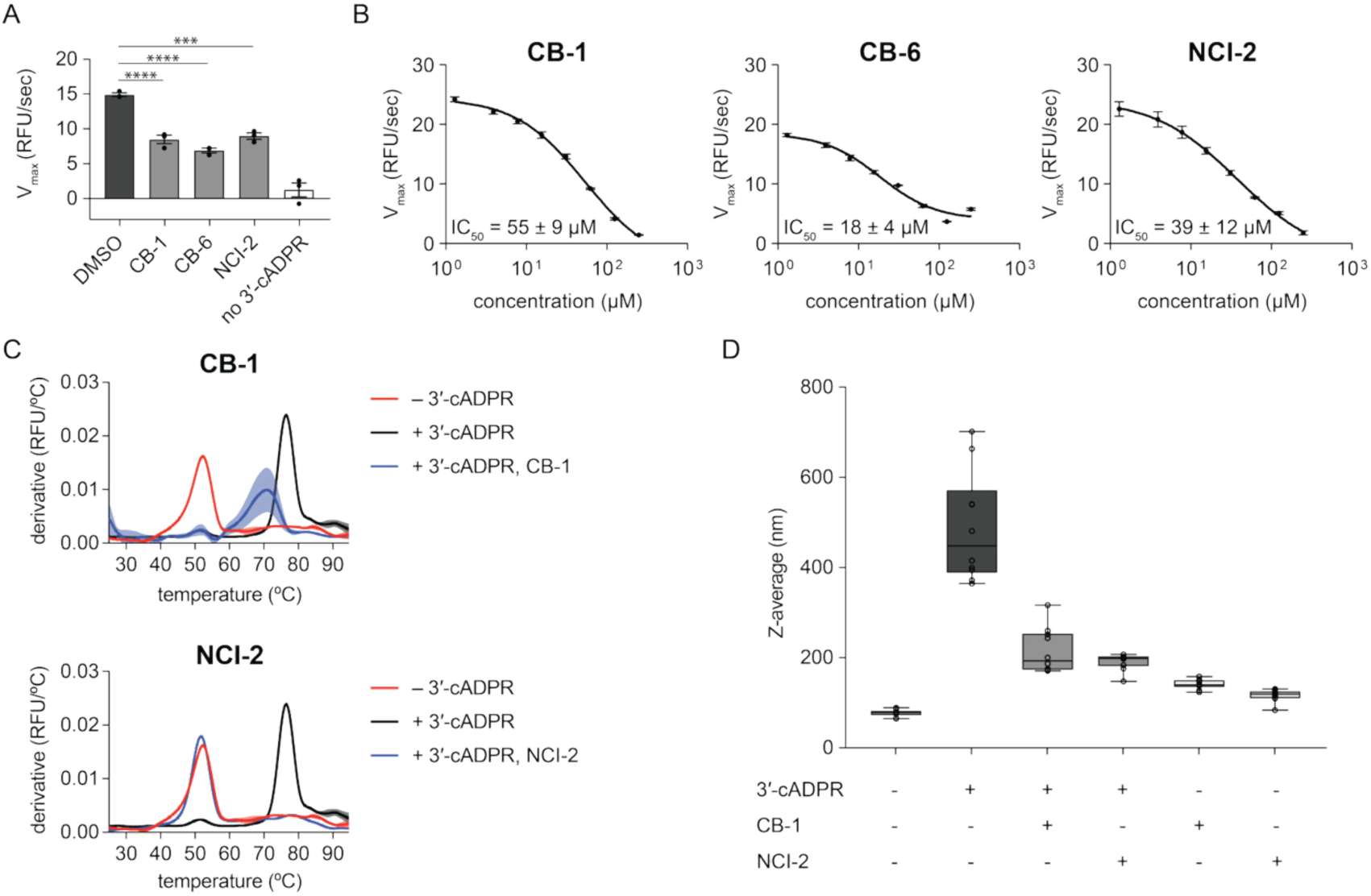
In vitro analyses reveal that the inhibitors prevent ThsA oligomerization and activation. A. Fluorescence induction by the hydrolysis of εNAD^+^ by ThsA (35 nM), activated by 3′-cADPR (35 nM) in the presence and absence of inhibitors (50 µM), displayed as mean ± SEM of independent experiments (n = 3). Results of an ordinary one-way ANOVA with a Dunnett’s multiple comparisons test using a single pooled variance are displayed (adjusted p values: **** p<0.0001, *** p=0.001). B. Dose-response curves of 3 inhibitors with IC_50_s ± SD from independent experiments (n = 3). C. Melting temperature-based quantification of 3′-cADPR-induced ThsA filamentation (0.45 µM of both ThsA and 3′-cADPR) in the presence and absence of inhibitors (1 mM each). Shaded error ranges represent SEM of independent experiments (n = 3). D. Dynamic light scattering quantification of 3′-cADPR-induced ThsA filamentation (0.45 µM of both ThsA and 3′-cADPR) in the presence and absence of inhibitors (500 µM each). Boxplots display each sample’s median and interquartile range. Whiskers extend to the minimum and maximum values. Each individual measurement (n = 10) is displayed as a circle.

Having established that the inhibitors target ThsA, we asked if they were directly inhibiting the NADase catalytic mechanism or acting upstream to inhibit the allosteric activation of ThsA. Previous work reported that ThsA is activated by a signal-induced filamentation process. ^21^ Therefore, we tested if the inhibitors prevented signal-induced filamentation. We tested this both by protein melting temperature analysis (3′-cADPR binding and filamentation increases T_m,_ Fig. 4C) and by dynamic light scattering (filamentation increases scattering, Fig. 4D). In both cases, the 3′-cADPR signal induced filamentation of ThsA, and both inhibitors prevented it. Neither inhibitor influenced filamentation on its own in the absence of 3′-cADPR signal (Fig. 4D). Therefore, both inhibitors CB-1 and NCI-2 appear to prevent the 3′-cADPR-induced filamentation process that activates ThsA.

To determine potential binding sites, we modeled the ThsA SLOG domain with the inhibitors using both Boltz-2 and MolModa. Using both methods, we saw that CB-1 and NCI-2 reside in the 3′-cADPR-binding pocket, suggesting competitive inhibition (Extended Data Fig. 3); however, experimental confirmation awaits the results of co-crystalization studies. Overall, our analyses demonstrate that type I Thoeris can be inhibited by targetting the signal-induced filamentation of ThsA. This finding also validates the filamentation-activation model of ThsA. ^21^

### Thoeris inhibitors allow low levels of immune-suppressed phages to overcome bacterial populations

Next we aimed to use these chemical inhibitors to address an outstanding question about antiviral immunity in bacteria. Namely, is an abortive infection system like Thoeris only required to eradicate new phages, or does it serve to continuously repress the same population of phages to low titers and/or dormant states? Both answers to this fundamental question have examples in the better-studied context of human antiviral immunity. Namely, human immune systems typically eradicate some viruses (e.g., influenza, hepatitis A), yet other viruses require continuous suppression by components of our immune system (e.g., hepatitis B and C). ^25,26^

To probe this question, we added the Thoeris-targeted SPO1 phage to a Thoeris-defended population of *B. subtilis*. We then waited several hours to supplement the culture with inhibitor CB-1. If the immune system can initially eradiate infectious phages, then the bacterial population would not succumb to phages, even after suppression of Thoeris immunity. However, if the immune system must constantly work to minimize phage replication (allowing low levels of phages to persist), then addition of a Thoeris inhibitor would shift the balance and allow the phages to overtake the bacterial population (Fig. 5A).

**Figure 5.**
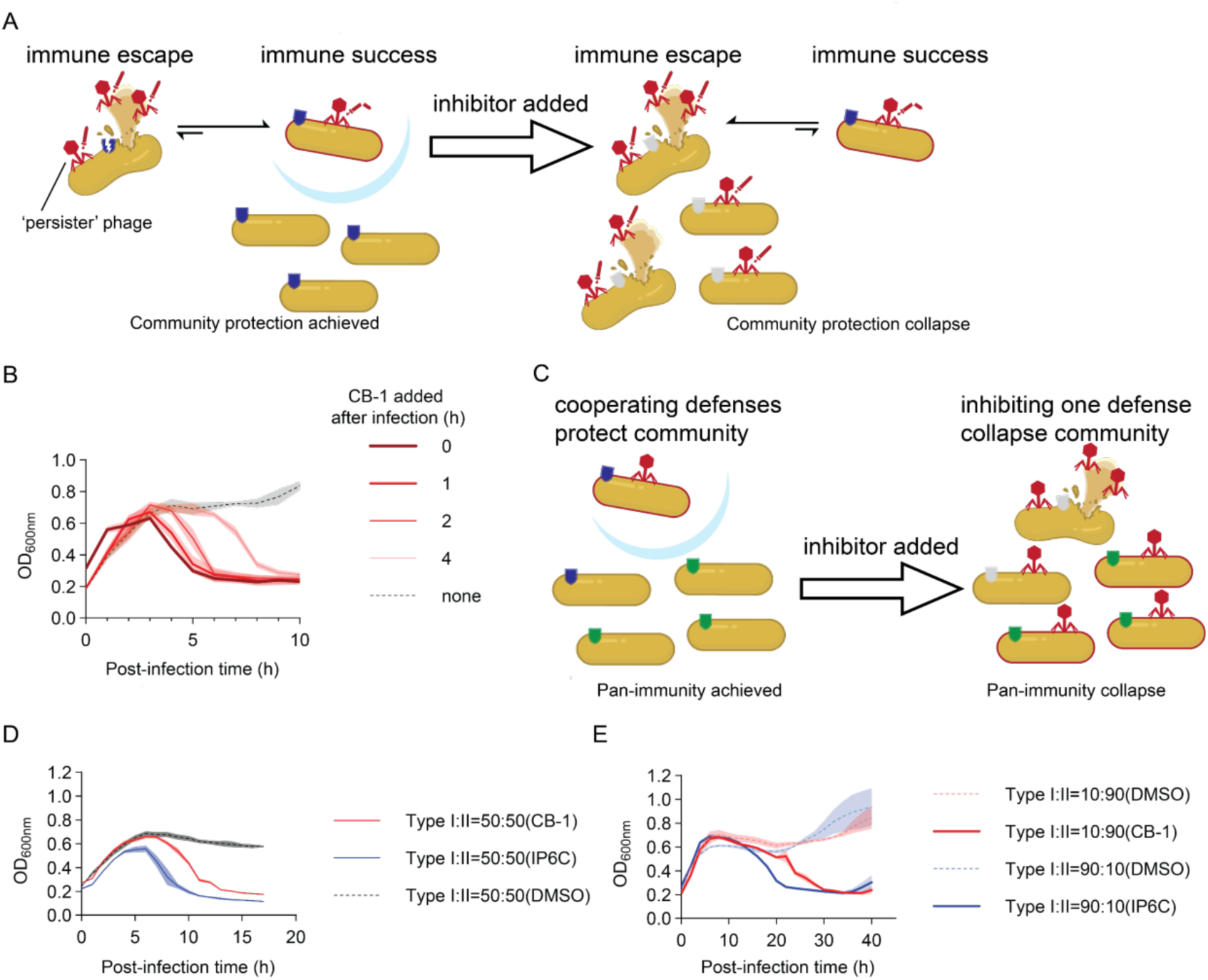
Thoeris inhibitors reveal that persistent and uniform antiviral immunity is necessary for phage defense. A: Schematic showing the hypothetical requirement for consistent antiphage immunity to actively repress low levels of phages. Immune suppression would revive ‘persister’ phages. B: Lysis curves of type I Thoeris-expressing *B. subtilis* infected with SPO1 phage (MOI = 0.01), followed by addition of inhibitor CB-1 (100 µM) after different time delays. See also Extended Data Fig. 4. C. Schematic showing the hypothetical population lysis resulting from inhibiting the immune system of only some bacteria in a cooperating community. D. Lysis curves of a 1:1 mixture of type I Thoeris-expressing *B. subtilis* and type II Thoeris-expressing *B. subtilis* infected with SPO1 phage (MOI = 0.01) and treated with either CB-1 (type I Thoeris inhibitor, 100 µM), IP6C (type II Thoeris inhibitor, 300 µM), or neither (DMSO). E. Lysis curves of a 1:9 and 9:1 mixtures of type I Thoeris-expressing *B. subtilis* and type II Thoeris-expressing *B. subtilis* infected with SPO1 phage (MOI = 0.01) and treated with either CB-1 (type I Thoeris inhibitor, 100 µM), IP6C (type II Thoeris inhibitor, 300 µM), or neither (DMSO). In panels B, D, and E, shaded error ranges represent SEM of a biological triplicate.

In all cases, delayed addition of CB-1 enabled SPO1 to overcome the bacterial population (Fig. 5B). We note that phage SPO1 contains the Tad2 protein, which helps it to partially evade type I Thoeris. ^27^ This counter-defense may make SPO1 uniquely suited to persist in the face of immunity and subsequently revive upon immune suppression. Therefore, we considered the possibility that the revival of persisting phages was unique to SPO1 (and other phages with mechanisms of partially circumventing abortive infection). However, we found that the two other phages we tested (SP50 and Goe2) yielded similar results upon delayed addition of CB-1 (Extended Data Fig. 4). Although SP50 contains the related Tad1 protein, ^17^ no known Tad proteins (Tad1–Tad6) were found in the Goe2 genome. ^28^ Therefore, low levels of persisting phages that evaded abortive infection can indeed overcome the host if the immune system is suppressed. Consequently, persistent viral suppression is a requirement for Thoeris.

Beyond fundamental biology, this question has implications for phage therapy. In prior work, we showed that simultaneous addition of lytic phages and an immune system inhibitor enabled clearance of bacterial populations—including pathogens in an animal infection model and single species within a mixed microbial model. ^15^ However, given our new result suggesting that low levels of infectious phages persist in the presence of antiphage immune systems, the simple addition of immune system inhibitors may actually revive endogenous phages to kill targeted bacteria without the need for delivering new phages.

### Inhibition of abortive infection in only 10% of cells collapses the entire bacterial population

Natural microbial environments often contain many bacterial species and even multiple strains within each species. ^29,30^ We still have only a nascent understanding of the significance of anti-phage immune systems in maintaining the balance of microbes in these environments. To fill this gap, chemical inhibitors of anti-phage immune systems may prove vital tools for ‘turning off’ individual immune systems and monitoring changes in microbial community structure.

We previously found that chemical inhibition of a type II Thoeris immune system could selectively deplete Thoeris-containing *P. aeruginosa* from a simple polymicrobial model containing only *P. aeruginosa*, *Staphylococcus aureus*, *Streptococcus sanguinis*, and the *P. aeruginosa* phage Lit1. ^15^ In this case, replication of phage Lit1 had no impact on the *S. aureus* and *S. sanguinis* bacteria because it selectively infects *P. aeruginosa*. However, in other cases, replication of a phage may deplete bacteria related to the immunity-depleted strain. Specifically, in environments where multiple strains of the same bacterial species co-exist, these strains may be sensitive to the same phage but each harbor their own immune systems to protect themselves. In this context, the bacterial strains also protect each other via ‘herd immunity’ by keeping the levels of their common viruses low. We hypothesized that inhibiting the abortive infection system of one strain may allow the common phage to propagate and deplete both strains from the population (Fig. 5C).

To test this hypothesis, we added the SPO1 phage to a mixture of two strains of *B. subtilis* that were identical except one possessed the type I Thoeris immune system discussed in this manuscript and the other possessed a type II Thoeris immune system from *Bacillus amyloliquefaciens* strain Y2. Both immune systems are abortive infection systems that can protect *B. subtilis* from SPO1. ^8^ As we described in this work, CB-1 inhibits the type I Thoeris, and in our previous work, we found an inhibitor of the type II Thoeris called IP6C. ^15^ We asked if addition of either CB-1 or IP6C alone could allow the SPO1 phage to deplete the entire population of both bacteria. Indeed, we found that either inhibitor (on its own) was sufficient to allow SPO1 to overcome the entire bacterial population if the two bacteria were mixed in a 1:1 ratio (Fig. 5D).

Previous research applying epidemiological models to understand the efficacy of abortive infection systems for ‘herd immunity’ showed that >90% (sometimes even 99%) of the bacteria must possess the defense to protect the population. ^31–34^ Therefore, we hypothesized that inhibiting the immune system of only 10% of the bacteria would enable the phage to kill the population. We tested both 1:9 and 9:1 mixtures of type I and type II Thoeris-defended strains, each treated only with the inhibitor of the minor represented strain. We found that when the immune system of only 10% of the bacteria was inhibited, the entire population was eventually lysed (Fig. 5E). Therefore, inhibiting an anti-phage immune system not only depletes the bacteria containing that immune system from a population, but it can also deplete other bacteria in the population that are targeted by the same phages and therefore rely on ‘herd immunity’ provided by the inhibited immune system.

## Discussion

We have discovered a set of selective chemical inhibitors of the type I Thoeris anti-phage immune system. Via biochemical characterization, we demonstrated that these inhibitors prevent the filamentation and activation of the ThsA effector NADase. This mechanism is reminiscent of SLOG domain inhibitors that inactivate the ADPR-activated TRPM2 channel in eukaryotes, ^35,36^ and it may apply broadly to other SLOG domain-containing antiviral defenses. In fact, since abortive infection systems generally involve effectors that are activated by alarm signals and/or filamentation, this strategy of chemical interference may prove broadly applicable across most abortive infection immune systems.

Additionally, due to their easy addition (and removal) at different time points, chemical probes afford the unique opportunity to dynamically assess the importance of immune systems over a time course. This allowed us to investigate the importance of *sustained* abortive infection immunity for antiviral defense. Our chemical inhibitor revealed that low levels of infectious phages remaining in “immune” bacterial populations can overcome the population after the immunity is lost. Although mechanistic differences abound between virus-host interactions in bacteria and humans, this finding is conceptually similar to how some human viruses can persist as minimally replicating viruses that can revive and overtake the host if immunity is suppressed (e.g., polyoma viruses and hepatitis B and C viruses). ^22,23,25,37^ Alternatively, based on the behavior of other myophages, SPO1 might enter a pseudolysogenic state^38–42^ that may evade Thoeris, which is conceptually related to latent human viruses like herpes viruses and human papillomavirus (HPV) that can also be activated upon immune suppression. ^43–48^ In either case, this finding suggests that there can be low levels of immune-targeted phages in populations with their potential hosts. Therefore, chemical inhibitors may be used on their own to shift microbial populations by de-repressing those endogenous immune-targeted phages. Phage therapies may even be possible by simply reviving repressed phages instead of requiring the addition of exogenous phages.

Furthermore, we found that the cooperative ‘herd immunity’ afforded by antiphage defenses can dramatically shape the population-level impact of inhibiting antiviral immunity. Related strains within a bacterial population rely on each others’ immunity for protection. If a population contains a mixture of strains of a species (and all are susceptible to a shared set of phages), then the entire mixture can collapse if an immune system is suppressed in only one of the strains. In this sense, a diversity of abortive infection immune systems across a population may be a liability, since it provides multiple routes for collapse. This liability to bacterial populations may partially explain the seemingly redundant expression of many immune systems within bacteria.

This finding also gives promise to the application of immune system inhibitors. Namely, the cooperative reliance on anti-phage immunity can compound the impact that even a weak inhibitor can have on a population. To be highly effective at promoting phage infection, a inhibitor of Thoeris (or other abortive infection systems) only needs to prevent abortive infection in 10% of the bacteria. In a therapeutic context, this is a remarkable advantage over antibiotics, which require universal high efficacy in all cells. Another implication of this ‘herd immunity’ is that low levels of spontaneous mutants resistant to inhibitor would still ultimately get killed by phages. Therefore, in contrast to antibiotics, resistance should spread slowly to inhibitors of abortive infection systems.

Although these type I Thoeris inhibitors were sufficiently potent and selective for our use, future medicinal chemistry efforts can likely improve these properties building off our scaffold. Furthermore, these inhibitors (and possibly more potent analogs) await testing against natural isolates with endogenous type I Thoeris systems. Based on our previous work with type II Thoeris inhibition, we expect this to largely succeed with systems that have similar ThsA proteins, but the potencies of the inhibitors may differ in ThsA homologs that are more divergent. ^19^

Beyond type I and type II Thoeris, over 100 antiphage immune systems exist, and it is still unclear how important most of them are in shaping microbial ecosystems and defending pathogens against phage therapies. ^5^ We expect a panel of chemical inhibitors against a diverse array of abundant immune systems would help fill these knowledge gaps. Given that our screen of only 1,581 compounds yielded a reliable inhibitor, we are optimistic that only minimal upfront effort may be required to discover selective chemical inhibitors of many of the remaining bacterial immune systems.

In conclusion, the type I Thoeris immune system can now be inhibited by compounds that selectively interfere with ligand-induced filamentation of ThsA. Our results confirmed the importance of filamentation for ThsA activity. Furthermore, we found that chemical depletion of bacterial immune systems can revive low levels of repressed phages to clear a bacterial population—an observation with both intriguing parallels to persistent human viral infections and implications for phage therapy. Finally, we found that herd immunity may cause entire mixed-strain communities to succumb to inhibitors that only suppress immunity in a small fraction of the bacteria. Ultimately these findings provide insight into the mechanism and functional importance of Thoeris immunity, and they provide optimism for the feasibility of developing a chemical toolkit to control the vast repertoire of antiviral immune systems in bacteria.

## Methods

### Construction of *B. subtilis* that carry the type I Thoeris system

The Thoeris cassette with its native promoter from *B. cereus* MSX-D12 [NCBI accession #AHEQ01000050, 16453-19685 (+)] was synthesized and cloned into the HindIII site on plasmid pDG1662 by GenScript. The constructed plasmids were propagated in *E. coli* DH5α and selected by 100 μg/mL ampicillin. The plasmids were extracted from *E. coli* DH5a using a QIAprep Spin Miniprep Kit. Then, the plasmids were transformed into *B. subtilis* RM125 using a protocol adapted from Young et al., ^49^ and a ‘no defense’ control strain was generated using the pDG1662 empty vector. Briefly, *B. subtilis* RM125 were grown in Medium A (1 g/L yeast extract, 0.2 g/L Casamino acids, 0.5% (w/v) glucose, 15 mM (NH_4_)_2_SO_4_, 80 mM K_2_HPO_4_, 44 mM KH_2_PO_4_, 3.9 mM sodium citrate, 0.8 mM MgSO_4_) at 37 °C, 220 rpm until the cessation of logarithmic growth and were allowed to continue growing for another 90 mins. Then, *B. subtilis* cells were diluted 10-fold into Medium B (Medium A + 0.5 mM CaCl_2_ + 2.5 mM MgCl_2_) and incubated at 37 °C, 300 rpm for 90 mins. 1 μg of plasmid was added to these now-competent cells and incubated at 37 °C, 220 rpm for 30 mins. Transformed *B. subtilis* cells with the double-crossover insertion at the amyE locus were selected based on resistance to 5 μg/mL chloramphenicol and sensitivity to 100 μg/mL spectinomycin.

### Defense protection assay on agar plates

Overnight cultures of *B. subtilis* RM125 transformed with type I Thoeris and empty pDG1662 vector were diluted 1:10 diluted into LB media. The diluted culture was poured onto 1.5% agar LB plates and left to absorb for 10 mins. After culture was removed, the plate was left to dry in a biological safety cabinet (∼30 min). After drying, 10-fold serial dilutions of SPO1, SP50, and Goe2 phage lysates were spotted onto the agar plates (5 µL each). The plates were incubated overnight at 37 °C.

### High throughput screening

A synthetic compound library from ChemBridge (DIVERSet-CL Library Block 1) and from the National Cancer Institute (NCI Diversity Library Set VII) were each used for screening. The compounds from the libraries were added to 384-well microtiter plates as 10 µL aliquots of 40 µM in LB + 1% DMSO. An overnight culture of *B. subtilis* type I (MSX-D12) Thoeris was diluted 1:100 into fresh LB media + 100 µg/mL spectinomycin and incubated at 37 °C, 220 rpm for 1 hour. Then, 20 µL of this freshly grown *B. subtilis* culture was added into each well of the 384-well plate (already containing 10 µL of inhibitor in each well), followed by incubation at 37 °C for 1 hour. Then, for the ChemBridge library, 10 µL of a 1:1:1 mixture of SPO1:SP50:Goe2 phages in LB (MOI = 0.01) was added to each well. For the NCI library screen, 10 µL of only SPO1 phage in LB (MOI = 0.01) was added to each well. The plates were incubated at 37 °C in a Biospa8 (Biotek), and the OD600_nm_ in each well was recorded every 1 hour using a Synergy H1 plate reader (Biotek).

### Hit validation experiments

An overnight culture of *B. subtilis* type I Thoeris was 1:50 diluted in LB media and incubated at 37 °C for ∼ 1h until its OD600_nm_ reached 0.1. 180 μL of this bacteria culture was added into each well of a 96-well plate containing a 3-fold serial dilution of the potential hit compounds (1000 µM – 0.15 µM). 20 μL of SPO1 phage stock (MOI = 0.01) was added to the wells and mixed by pipetting. The plates were incubated at 37 °C in a Biospa8 (Biotek), and the OD600_nm_ in each well was recorded every 1 hour using a Synergy H1 plate reader (Biotek). The concentration with the strongest lysis profile was reported in Extended Data Figure 1B&C.

### Spot-on-lawn antimicrobial experiments

An overnight culture of *B. subtilis* RM125 transformed with type I Thoeris was diluted 1:25 into 5 mL 0.5% agar LB media. This soft agar culture was poured onto 1.5% agar LB plates and left to solidify. After cooling, 3-fold serial dilutions of each potential inhibitor in a DMSO solution were spotted onto the agar plates (2 µL each, 10 mM –123µM). The plates were incubated overnight at 37 °C.

### Construction of *B. subtilis* that only expresses MSX-D12 ThsB

MSX-D12 ThsB was amplified by PCR using pDG1662: type I Thoeris as the template with primers ThsB_pDR110_F and ThsB_pDR110_R, followed by DpnI digestion and PCR cleanup using QIAquick PCR Purification Kit (QIAGEN #28104). pDR110 was amplified by PCR using primers pDR110_F and pDR110_R, followed by DpnI digestion and gel purification using QIAquick Gel Extraction Kit (QIAGEN #28704). MSX-D12 ThsB was ligated with pDR110 using NEBuilder® HiFi DNA Assembly Master Mix (New England Biolab #E2621) and electroporated into NEB® 10-beta Electrocompetent *E. coli* (New England Biolab #C3020K), which was selected by 100 µg/mL ampicillin. The sequence of the constructed plasmid was verified by whole plasmid sequencing. MSX-D12 ThsB was placed downstream of the P*_spank_* promoter (IPTG-inducible) on plasmid pDR110. The *B. subtilis* RM125 was transformed with pDR110 plasmid using the transformation method above.

### Preparation of phage-infected cell lysate for LC-HRMS analysis

Overnight cultures of *B. subtilis* Pspank negative control or *B. subtilis* P_spank_ MSX-D12 ThsB were diluted 1:100 into 500 mL fresh LB + 100 μg/mL spectinomycin + 1 mM IPTG. When testing inhibitors, 1 mM of the compound was supplemented to the *B. subtilis* P_spank_ MSX-D12 ThsB culture. The diluted cultures were incubated at 30 °C, 220 rpm for 4 hours until OD_600nm_∼0.3. 50 mL of the culture was removed as a t=0 min sample and immediately centrifuged at 10,000 × g at 4 °C for 5 mins. The supernatant was discarded, and the cell pellet was stored at −80 °C. Then, 5 mL of SPO1 phage (∼5×10^10^ PFUs/mL) was added to the host cells to reach MOI∼10. The infected cell culture was incubated at 30 °C, 220 rpm, and 50 mL of the culture was removed at different time points to be immediately centrifuged at 10,000 × g, 4 °C for 5 mins. The supernatant was discarded, and the cell pellet was stored at −80 °C. The cell pellets were thawed at room temperature and resuspended in 600 μL of 100 mM sodium phosphate buffer (pH = 7) + 4 mg/mL lysozyme. After incubation at room temperature for 10 mins, the cells were transferred into 2 ml tubes with Lysing Matrix B and lysed using an Omni Bead Ruptor 12 for 2 × 40 s at 6 m/s with a dwell time of 4 mins in-between. After lysis, the tubes were centrifuged at 15,000 × g at 4 °C for 10 mins. Then, 400 μL of each supernatant were transferred to Amicon Ultra-0.5 Centrifugal Filter Units 3 kDa and centrifuged for 45 mins at 14,000 g 4 °C. The filtrate was collected, and 10 μl of each were used for LC-MS analysis.

### LC-MS analysis of 3′-cADPR, and NAD^+^

The liquid chromatography analysis was performed on ACQUITY UPLC I-Class PLUS System using a Luna Omega 5 μm Polar C18 100 Å column (250×4.6 mm). The mobile phase A was water + 0.1 % (v/v) formic acid and the mobile phase B was acetonitrile + 0.1 % (v/v) formic acid. The flow rate was kept at 0.7 mL·min^−1^ and the gradient was as follows: 0% B (0–10 min), increase to 2.5% B (10–15 min), increase to 5% B for (15–16 min), 0% B (16–26 min), increase to 95% B (26–27 min), 100% B (27–37 min), decrease to 0% B (37–38 min), 0% B (38–48 min). High-resolution electrospray ionization (HR-ESI) mass spectra with collision-induced dissociation (CID) MS/MS were obtained using a Waters Synapt G2S Quadrupole Time-of-Flight (QTOF). The instrument was operated at negative ionization mode. The MS spectra were obtained on Quadrupole analyzer with a scan range of 300–800 Da and analyzed using MassLynx 4.1 software. The *m/z* of interest from Quadrupole was subjected to CID (energy ramp 34–44 V) and analyzed on the Time-of-Flight analyzer with a scan range of 50–750 Da. “Normalized 3′-cADPR intensity” was calculated by dividing the 3′-cADPR intensity by the NAD^+^ intensity for that same lysate, followed by dividing this quotient by the mean of the quotients for the “no inhibitor” control samples. Therefore, a compound that fails to inhibit 3′-cADPR production would have a value ∼ 1.

### Testing in vitro NADase activity of ThsA with fluorogenic εNAD^+^ substrate

ThsA was expressed and purified as described previously. ^21^ Enzymatic hydrolysis of the εNAD nicotinamide glycosidic bond causes the product εADPR to produce a fluorescent signal. Reactions (60 μl) in reaction buffer (10 mM HEPES pH 7.5, 150 mM NaCl, 5 mM MgCl_2_) containing 100 μM εNAD^+^ (Sigma #N2630), 35 nM ThsA, 35 nM 3′-cADPR (Biolog #C404), and inhibitor (1.3-250 µM) were prepared in a 96-well transparent flat-bottom plate (Greiner catalogue No. 655101) and loaded into a CLARIOstar Plus (BMG LABTECH) plate reader, in which fluorescence was measured using 300 nm excitation and 410 nm emission wavelengths in intervals of 30 s using orbital scanning and averaging of eight measurements at 25 °C. The maximum reaction rate (in RFU/seconds) was determined using linear regression on the linear part of the initial reaction curve in R (v4.0) software. ^50^

### Testing in vitro filamentation of ThsA by melting temperature quantification

Nano differential scanning fluorimetry (nanoDSF) experiments were performed on a Prometheus NT.48 device (NanoTemper Technologies) using 0.45 μM of ThsA, 0.45 µM 3′-cADPR, and 500 µM inhibitor was prepared in reaction buffer.

### Testing in vitro filamentation of ThsA by dynamic light scattering

A 10 μL sample containing 0.45 μM of ThsA, 0.45 µM 3′-cADPR, and 500 µM inhibitor was prepared in reaction buffer. Sample was loaded into a 2 microlitre quartz cuvette (Malvern Panalytical) and measurements were carried out with a Zetasizer μV photometer (Malvern Panalytical) using Zetasizer Software (v6.20). Measurements were carried out in 1-min runs containing six 10-s measurements. Each run produced a z-average value, corresponding to the harmonic-intensity-averaged particle diameter, which was plotted over time. 10 replicate runs were performed for each condition.

### Docking of inhibitors into ThsA

MolModa 1.0.1^51^ was used to dock inhibitors CB-1 and NCB-2 to the type I ThsA SLOG domain (PDB ID 7UXS, chain A^19^). Prediction of CB-1 and NCB-2 bound SLOG domain structures (100 models) were performed with the Boltz2. ^52^ SMILES sequences of the inhibitors were obtained in OpenBabel 3.1.0^53^, cif and pdb files of the inhibitors were generated using AceDRG^54^ and SMILES as inputs. The figures were prepared using Pymol.

### Testing delayed addition of inhibitor CB-1 against Thoeris defense

An overnight LB culture of *B. subtilis* type I Thoeris was 1:50 diluted in LB media and incubated at 37 °C for ∼ 1h until OD600nm reached 0.1. 180 μL of this diluted bacterial culture was added to each well of a 96-well plates. 20 μL of SPO1/Goe2 phage stock (MOI = 0.01) or SP50 phage stock (MOI = 10^−4^) was added to the wells and mixed by pipetting. 100 μM inhibitor CB-1 was added at 0, 1, 2, or 4 hours after phage addition. The plates were incubated at 37 °C in a Biospa8 (Biotek), and the OD600nm in each well was recorded every 1 hour using a Synergy H1 plate reader (Biotek).

### Testing phage lysis promotion of inhibitors against mixed populations of type I and type II Thoeris-containing *B. subtilis*

Overnight LB cultures of *B. subtilis* expressing type I Thoeris (this study) and type II Thoeris (discovered previously^15^) were 1:50 diluted into fresh LB, incubated at 37 °C for ∼ 1h until OD600_nm_ reached 0.1. For 50:50 mixing, 90 μL of each type I and II Thoeris bacteria culture was added into each well of a 96-well plate containing 100 μM CB-1 or 300 μM IP6C inhibitor. For 1:9 mixing, 18 μL and 162 μL type I and II Thoeris bacteria culture was added into each well of a 96-well plate containing 100 μM CB-1 or 300 µM IP6C inhibitor. 20 μL of SPO1 phage stock (MOI = 0.01) was added to the wells and mixed by pipetting. The plates were incubated at 37 °C in a Biospa8 (Biotek), and the OD600_nm_ in each well was recorded every 1 hour using a Synergy H1 plate reader (Biotek).

## Data availability

The sequence of phage SP50 is available on NCBI (accession number in progress). All other data generated in this study is included in the manuscript and supporting information. Any further requests for raw data should be addressed to the corresponding author.

## Acknowledgements

We thank the National Cancer Institute Development Therapeutics Program (NCI/DTP) for providing compounds present in this manuscript, specifically, the Diversity Library Set VII and NCI-2 (NSC #: 131986). We thank Adam Zlotnick (Indiana University) for access to the Chembridge DIVERSet library and Jared Lewis (Indiana University) for liquid handling assistance. We thank Tuli Mukhopadhyay for helpful discussions. We thank the Bacillus Genomic Stock Center (Ohio State University) and Robert Hertel (University of Goettingen) for providing bacteria and phages. The research was supported by a National Science Foundation CAREER award (IOS-2143636) to J.P.G. and a Camille Dreyfus Teacher-Scholar Award (TC-24-028) to J.P.G. Research support was also provided by the Research Council of Lithuania (LMTLT) agreement no. S-MIP-24-84 (to G.T.).

## Author contributions

Conceptualization, C.Z., Y.C., Z.Z., J.P.G.; methodology, C.Z., D.S., Z.Z., G.T., J.P.G.; analysis, C.Z., D.S., Y.C., G.T., J.P.G.; investigation, C.Z., D.S., Y.C.; writing – original draft, C.Z.; writing – review & editing, C.Z., D.S., Y.C., Z.Z., G.T., J.P.G.; visualization, C.Z., D.S., G.T., J.P.G.; supervision, G.T., J.P.G.; funding acquisition, G.T., J.P.G.

## Materials & Correspondence

Correspondence and requests for materials should be addressed to Joseph P. Gerdt.

## Extended Data Figures

**Extended Data Figure 1.**
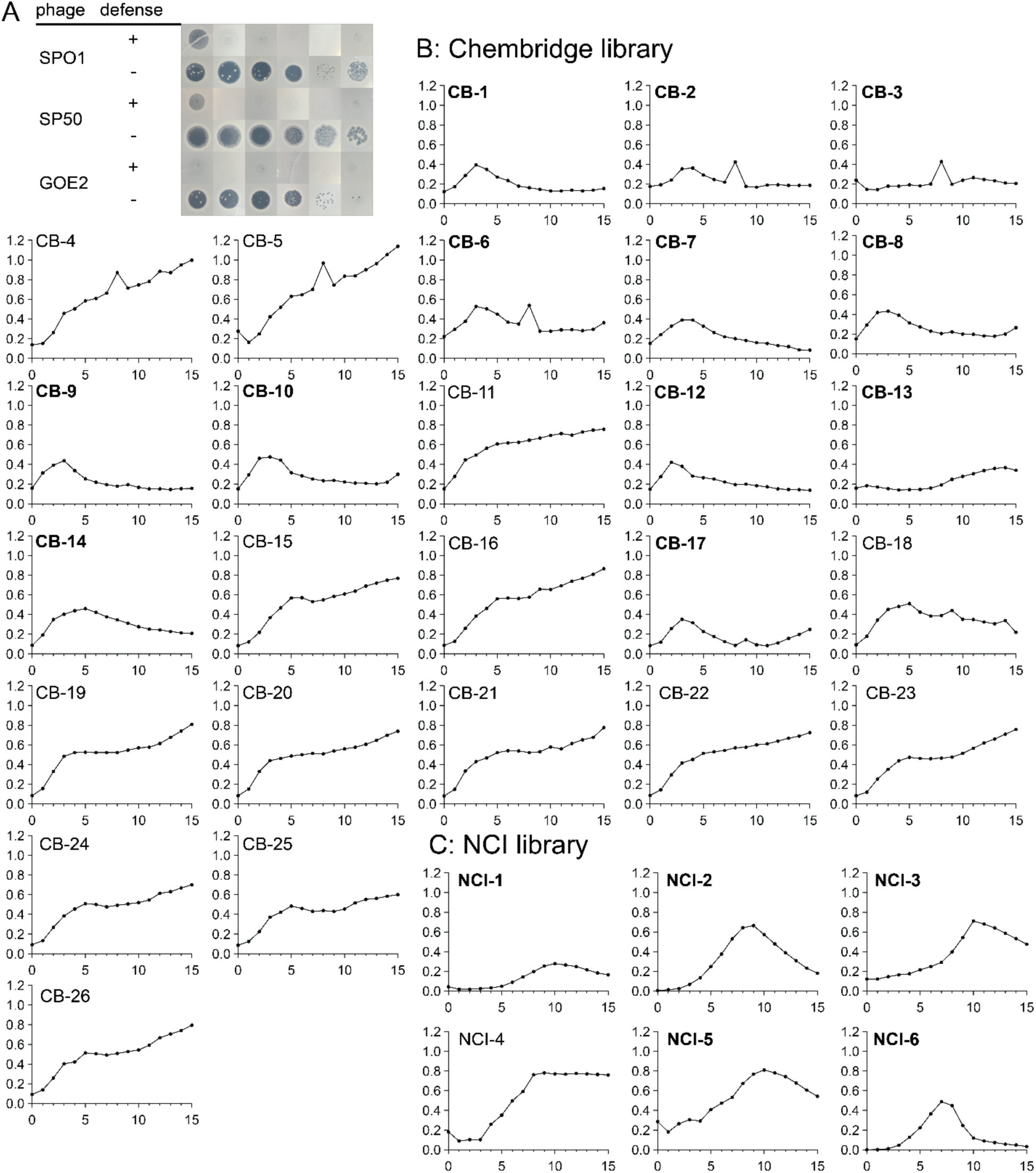
Type I Thoeris agar protection against 3 phages and validation of screening hits. A: Type I Thoeris protects *B. subtilis* against 3 phages. Images display a 10-fold serial dilution of each phage spotted onto a lawn of *B. subtilis* expressing or lacking the type I Thoeris defense. B–C. Validation of the screening hits by re-testing each at a range of concentrations (1000 µM – 0.15 µM) and monitoring for unambiguous culture lysis by phage SPO1 (MOI = 0.01). The treatment concentration with the clearest lysis is displayed here. Panel B is the Chembridge library; panel C is the NCI library. Bolded inhibitor codes are those that repeated lysis. X axis is post-infection time (hours). Y axis is OD_600nm_.

**Extended Data Figure 2.**
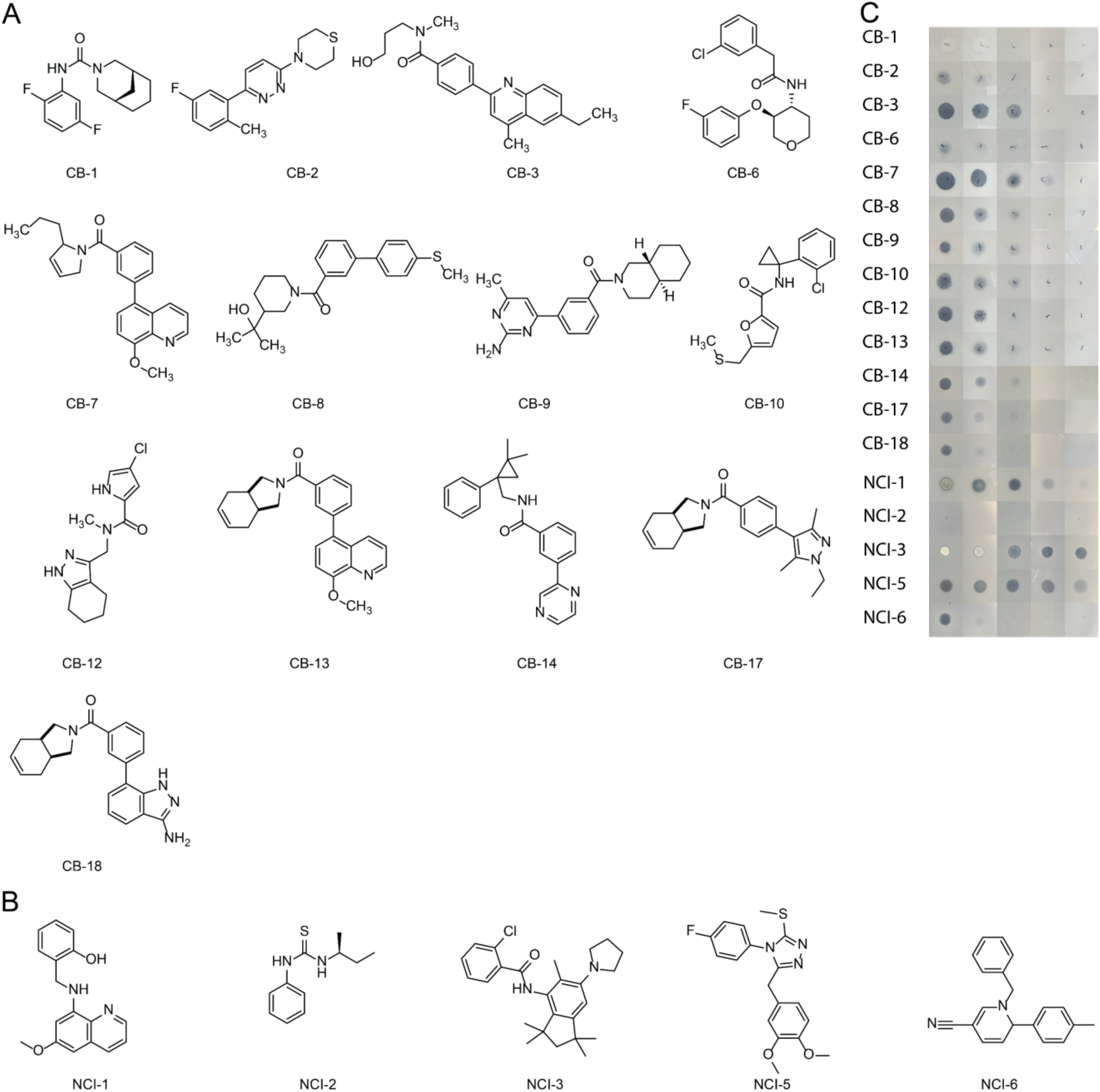
Structures of validated inhibitors and spot-on-lawn analysis for toxicity of inhibitors. A–B. Chemical structures of the inhibitors validated in Extended Data Fig. 1. C. Photos of spot-on-lawn antimicrobial assays testing the toxicity of the validated inhibitors. 2 µl of 3-fold serially diluted (10 mM –123µM) DMSO solutions of each inhibitor were spotted onto a lawn of *B. subtilis* without any phages. Zones of growth inhibition reveal non-selective antimicrobial activity.

**Extended Data Figure 3.**
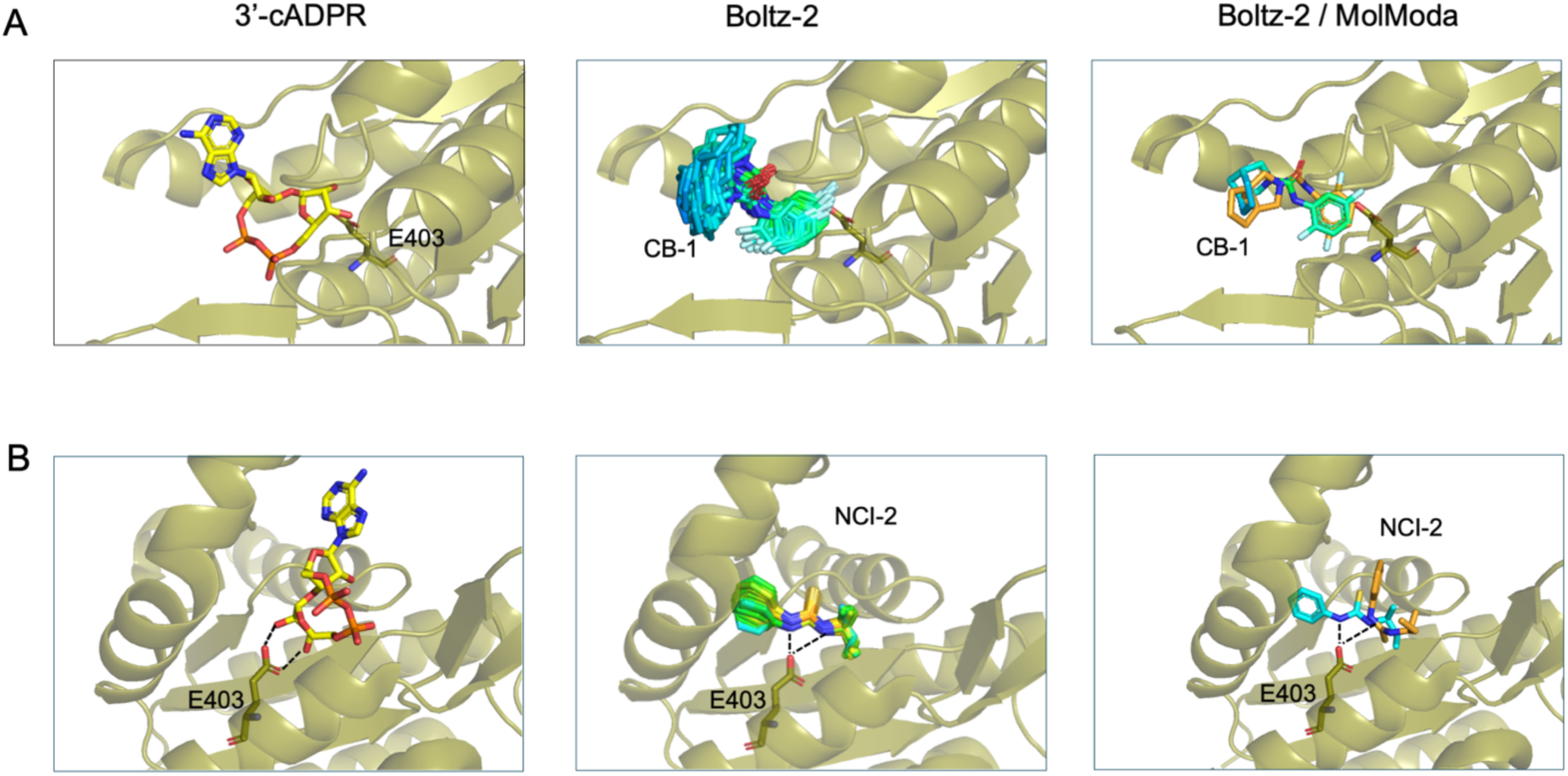
Predictions of inhibitors CB-1 and NCI-2 binding to the SLOG domain of ThsA. Comparison of the 3′-cADPR binding to the ThsA SLOG domain (PED ID 7UXS chain A^19^), left panel, to the Boltz-2 modeling largest cluster (middle panel) and Boltz-2 selected model from the largest cluster (cyan) with the best MolModa docking model (orange) (right panel). A. Prediction results for the CB-1 inhibitor, the largest cluster of 57 of 100 Boltz-2 models are shown (middle panel). B. Prediction of NCI-2 binding. The largest cluster of 84 of 100 models predicted by Boltz-2 is shown in the middle panel.

**Extended Data Figure 4.**
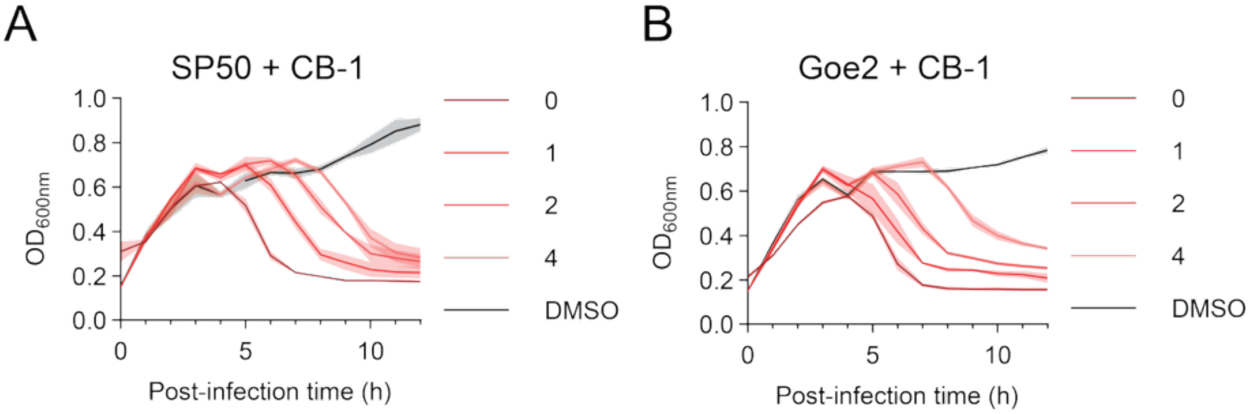
Phage persistence and revival upon immunosuppression in different infection-inhibition models. A–B. Lysis curves of type I Thoeris-expressing *B. subtilis* infected with SP50 (A, MOI = 0.01) or Goe2 phage (B, MOI = 0.0001), followed by addition of inhibitor CB-1 (100 µM) after different time delays.

## Supporting Information

### Supplementary tables

**Table S1.** Strains and bacteriophages used in this study.

| Bacterial strains | Description | Source |
| --- | --- | --- |
| *B. subtilis* RM125 | Wild type strain | Bacillus Genetic Stock Center #1A253 |
| *B. subtilis* pDG1662 | RM125 with pDG1662 inserted at the amyE locus on the genome. No Thoeris control. | This study |
| *B. subtilis* Y2 Thoeris | RM125 with pDG1662:Y2 Thoeris inserted at the amyE locus on the genome | Gerdt lab collection<sup>1</sup> |
| *B. subtilis* MSX-D12 Thoeris | RM125 with pDG1662:MSX-D12 Thoeris inserted at the amyE locus on the genome | This study |
| *B. subtilis* Pspank | RM125 with pDR110 inserted at the amyE locus on the genome. P<sub>spank</sub> promoter only control. | This study |
| *B. subtilis* Pspank-MSX-D12 ThsB | RM125 with pDR110:MSX-D12 ThsB inserted at the amyE locus on the genome. ThsB expression is controlled by an IPTG-inducible promoter (P<sub>spank</sub>) | This study |
| **Bacteriophages** |  |  |
| SPO1 | Bacillus phage | Bacillus Genetic Stock Center #1P4 |
| SP50 | Bacillus phage | The Félix D'Hérelle Reference Center for Bacterial Viruses HER 174 |
| Goe2 | Bacillus phage | Daniel Schwartz, Indiana University<sup>2</sup> |
| **Plasmids** |  |  |
| **pDG1662** | Empty vector that integrates at amyE locus on *Bacillus subtilis* genome | Bacillus Genetic Stock Center #ECE113 |
| **pDG1662:Y2 Thoeris** | Vector carrying Thoeris cassette from *B. amyloliquefaciens* Y2 that integrates at amyE locus on *Bacillus subtilis* genome | Gerdt lab collection<sup>1</sup> |
| **pDG1662:MSX-D12 Thoeris** | Vector carrying Thoeris cassette from *B. cereus* MSX-D12 that integrates at amyE locus on *Bacillus subtilis* genome | This study |
| **pDR110** | Empty vector carrying $P_{spank}$ promoter (IPTG-inducible) that integrates at amyE locus on *Bacillus subtilis* genome. | Bacillus Genetic Stock Center #ECE311 |
| **pDR110:MSX-D12 ThsB** | Vector carrying MSX-D12 ThsB which is downstream of $P_{spank}$ promoter that integrates at amyE locus on *Bacillus subtilis* genome. | This study |

**Table S2.** Chemical used in this study.

| **Chemicals** | **Source** | **Identifier** |
| --- | --- | --- |
| **LB broth** | VWR | 90003-350 |
| **Dehydrated Agar** | Fisher | DF0140-07-4 |
| **Agarose** | VWR | 0710 |
| **MgSO<sub>4</sub>** | Sigma Aldrich | M3634 |
| **Spectinomycin** | Sigma Aldrich | S9007 |
| **Ampicillin** | Sigma Aldrich | A9518 |
| **Chloramphenicol** | Sigma Aldrich | C0378 |
| **DMSO** | Sigma Aldrich | D8418 |
| **IPTG** | Gold Biotechnology | I2481C50 |
| **Formic acid (HPLC)** | VWR | PI85178 |
| **Sodium phosphate dibasic** | Sigma Aldrich | S5136 |
| **Sodium phosphate monobasic** | Sigma Aldrich | S3139 |
| **lysozyme** | Research Products International | L38100 |
| **DpnI** | New England Biolabs | CR0176 |
| **CB-1** | ChemBridge | 16969232 |
| **CB-2** | ChemBridge | 39134971 |
| **CB-3** | ChemBridge | 21496966 |
| **CB-4** | ChemBridge | 62784882 |
| **CB-5** | ChemBridge | 33206481 |
| **CB-6** | ChemBridge | 31767446 |
| **CB-7** | ChemBridge | 98700676 |
| **CB-8** | ChemBridge | 32633041 |
| **CB-9** | ChemBridge | 87683073 |
| **CB-10** | ChemBridge | 34348762 |
| **CB-11** | ChemBridge | 25169461 |
| **CB-12** | ChemBridge | 12066781 |
| **CB-13** | ChemBridge | 98410359 |
| **CB-14** | ChemBridge | 34368337 |
| **CB-15** | ChemBridge | 29385702 |
| **CB-16** | ChemBridge | 47497750 |
| **CB-17** | ChemBridge | 19054682 |
| **CB-18** | ChemBridge | 68914988 |
| **CB-19** | ChemBridge | 84634431 |
| **CB-20** | ChemBridge | 41121620 |
| **CB-21** | ChemBridge | 57074907 |
| **CB-22** | ChemBridge | 76489351 |
| **CB-23** | ChemBridge | 68248501 |
| **CB-24** | ChemBridge | 81061167 |
| **CB-25** | ChemBridge | 73783921 |
| **CB-26** | ChemBridge | 49155457 |
| **NCI-1** | NCI Diversity Set VII | 130801 |
| **NCI-2** | NCI Diversity Set VII | 131986 |
| **NCI-3** | NCI Diversity Set VII | 321496 |
| **NCI-4** | NCI Diversity Set VII | 201989 |
| **NCI-5** | NCI Diversity Set VII | 379388 |
| **NCI-6** | NCI Diversity Set VII | 351674 |

**Table S3.**
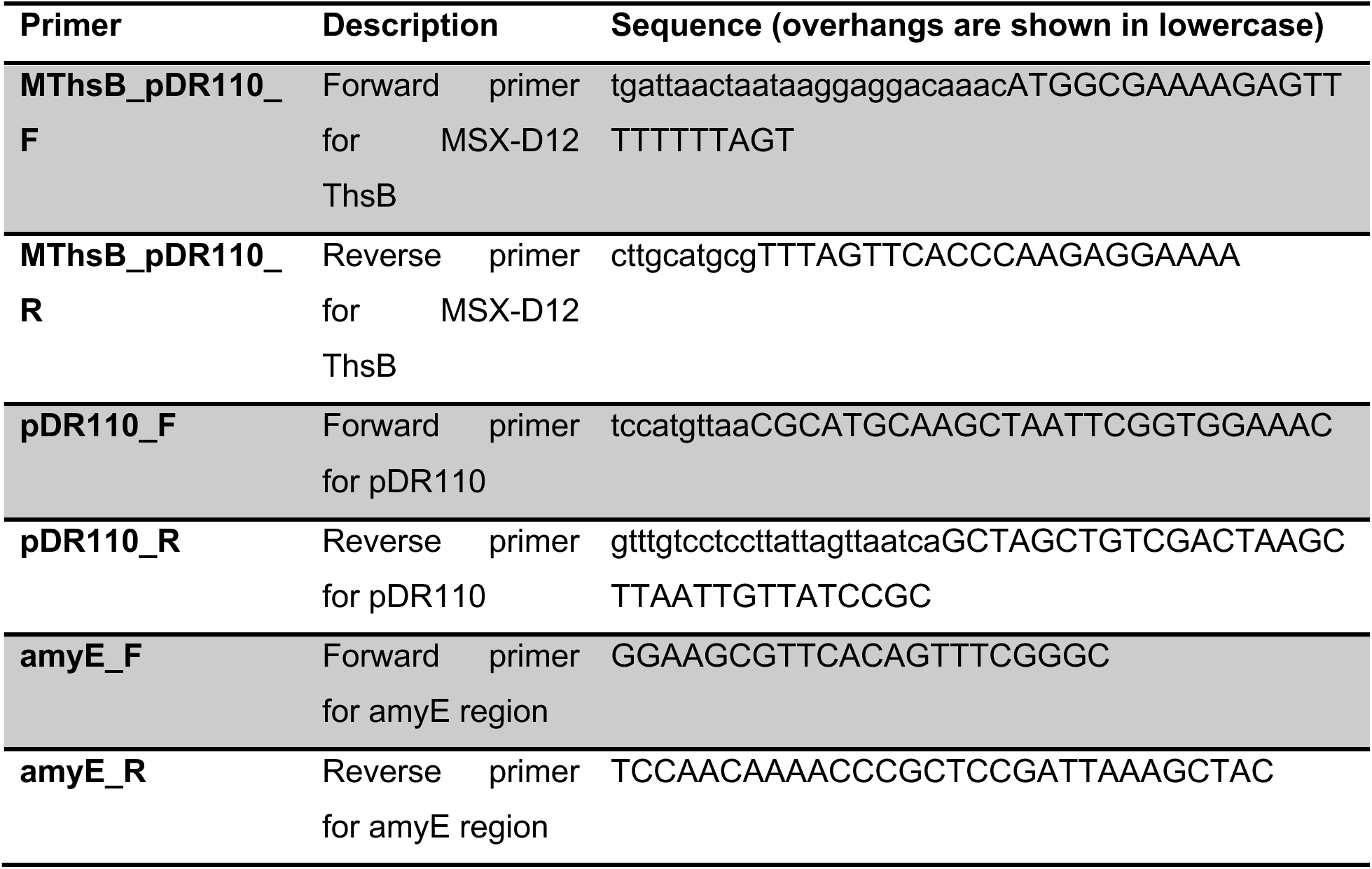
Primers used in this study.

